# Gut Microbial Butyrate Metabolism Generates an *N*-Acyl Amide that Regulates Host Transcriptional Programs

**DOI:** 10.64898/2026.11.16.766927

**Authors:** Randy Hamchand, Pamela V. Chang

**Affiliations:** Department of Microbiology and Immunology, Cornell University, Ithaca, New York, USA; Department of Chemistry and Chemical Biology, Cornell University, Ithaca, New York, USA; Cornell Center for Immunology, Cornell University, Ithaca, New York, USA; Cornell Institute of Host Microbe Interactions and Disease, Cornell University, Ithaca, New York, USA; Cornell Center for Innovative Proteomics, Cornell University, Ithaca, New York, USA

## Abstract

*Faecalibacterium prausnitzii* is a prominent member of the human gut microbiota whose depletion is associated with various intestinal and metabolic disorders. While multiple mechanisms have been proposed to explain how *F. prausnitzii* promotes host health, the molecular determinants underlying these activities remain elusive. Here, we show that *F. prausnitzii* is a prolific producer of the human metabolite *N*-butyrylglycine, which we find accumulates to millimolar concentrations in bacterial culture and is readily detectable in the feces of conventional wildtype mice. We establish an enzymatic basis of *N*-butyrylglycine production by identifying and biochemically validating an *F. prausnitzii N*-acyltransferase that catalyzes the conjugation of butyryl-CoA and glycine. Despite being derived from butyrate, *N*-butyrylglycine exhibits distinct bioactivity from its precursor, including reduced histone deacetylase inhibitory activity, an altered inflammatory profile, and enhanced suppression of cellular proliferation across multiple cancer cell lines. Furthermore, transcriptomic profiling revealed *N*-butyrylglycine coordinates repression of G2/M checkpoint-associated programs and downregulation of the colorectal cancer-associated gene *AOC1*. Together, these findings suggest that specialized metabolites produced by *F. prausnitzii* contribute to its beneficial effects and may underlie its protective role in colorectal cancer.

## Introduction

In the last decade, *Faecalibacterium prausnitzii* has emerged as one of the most important commensal bacteria of the human gut microbiota.^1,2^ Loss of *F. prausnitzii* has been consistently associated with intestinal and metabolic disorders including inflammatory bowel disease,^3,4^ type 2 diabetes,^5^ and colorectal cancer (CRC).^6,7^ Consequently, substantial effort has been invested towards understanding how *F. prausnitzii* influences host health. Much of this work has focused on the bacterium’s ability to produce butyrate,^8,9^ a short-chain fatty acid (SCFA) that regulates host physiology through mechanisms including histone deacetylase inhibition,^10^ epithelial barrier support,^11^ and cellular proliferation.^12^ However, butyrate production alone is unlikely to fully explain the organism-specific effects associated with *F. prausnitzii*. Butyrate biosynthesis is conserved among phylogenetically diverse gut anaerobes,^13–15^ and some butyrate-producing taxa, such as *Fusobacterium nucleatum*, are instead associated with CRC progression and poor clinical outcomes.^16^ These observations suggest that the biological activities of *F. prausnitzii* may extend from additional metabolic features.

## Results and Discussion

To investigate this possibility, we profiled the *F. prausnitzii* metabolome. Motivated by recent studies that linked select *F. prausnitzii* strains to fatty acid metabolism,^17^ we focused on carboxylic acid-containing metabolites. To enable sensitive detection of organic acids and related compounds, culture extracts were derivatized with 3-nitrophenylhydrazine (3-NPH) using an *N*-(3-dimethylaminopropyl)-*N*’-ethylcarbodiimide (EDC)-mediated coupling strategy and subjected to liquid chromatography-mass spectrometry (LC-MS) analysis. (**Fig. 1**).^18^

**Fig. 1.**
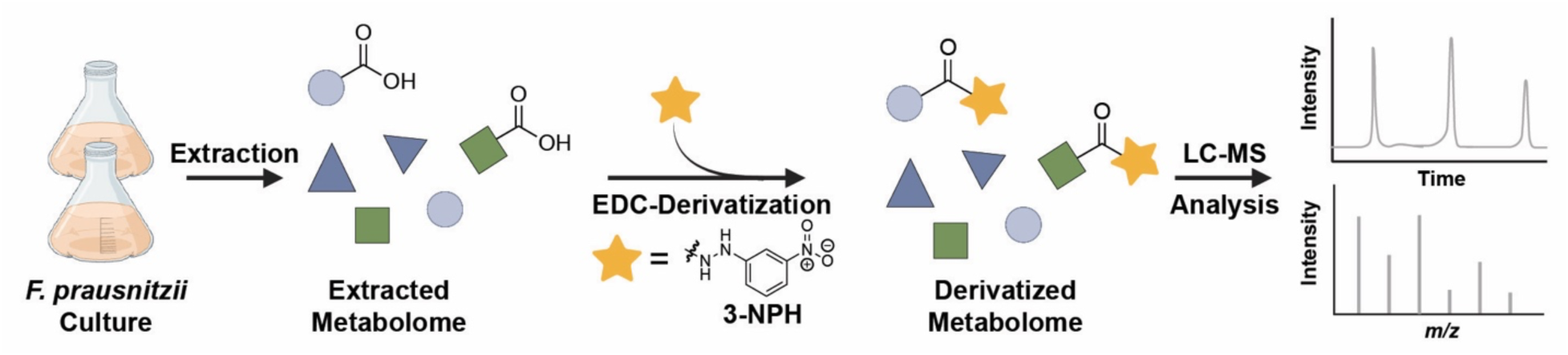
Workflow for EDC-mediated 3-nitrophenylhydrazine derivatization and LC-MS analysis of carboxylic acid-containing metabolites produced by *F. prausnitzii*. EDC = *N*-(3-dimethylaminopropyl)-*N*’-ethylcarbodiimide.

Consistent with literature, this analysis showed strong production of the SCFAs acetate and butyrate.^19^ Importantly, we also detected an additional prominent 3-NPH-derivatized metabolite with [M – H]^-^ = 279.1100 Da. Tandem MS analysis confirmed characteristic fragments arising from the 3-NPH derivatization group and revealed a neutral loss of 70.0433 Da, corresponding to the elemental composition C_4_H_6_O and consistent with cleavage of a butyryl moiety (**Fig. S1**). This fragmentation generated a product ion at *m/z* 209.0686, in agreement with the expected mass of 3-NPH-derivatized glycine ([M – H]^-^_theoretical_ = 209.0680 Da, [M – H]^-^_obs_ = 209.0686 Da). Together this fragmentation data is consistent with the *N*-acyl amino acid *N*-butyrylglycine (NButGly) (**Fig. 2a**). To verify this structural assignment, we synthesized NButGly and compared it to the feature detected in *F. prausnitzii* culture extracts (**Fig. S2-4**). The synthetic standard and natural product exhibited identical chromatographic retention times and MS^2^ fragmentation patterns, confirming that *F. prausnitzii* produces NButGly (**Fig. 2b**, **Fig. S5**).

**Fig. 2.**
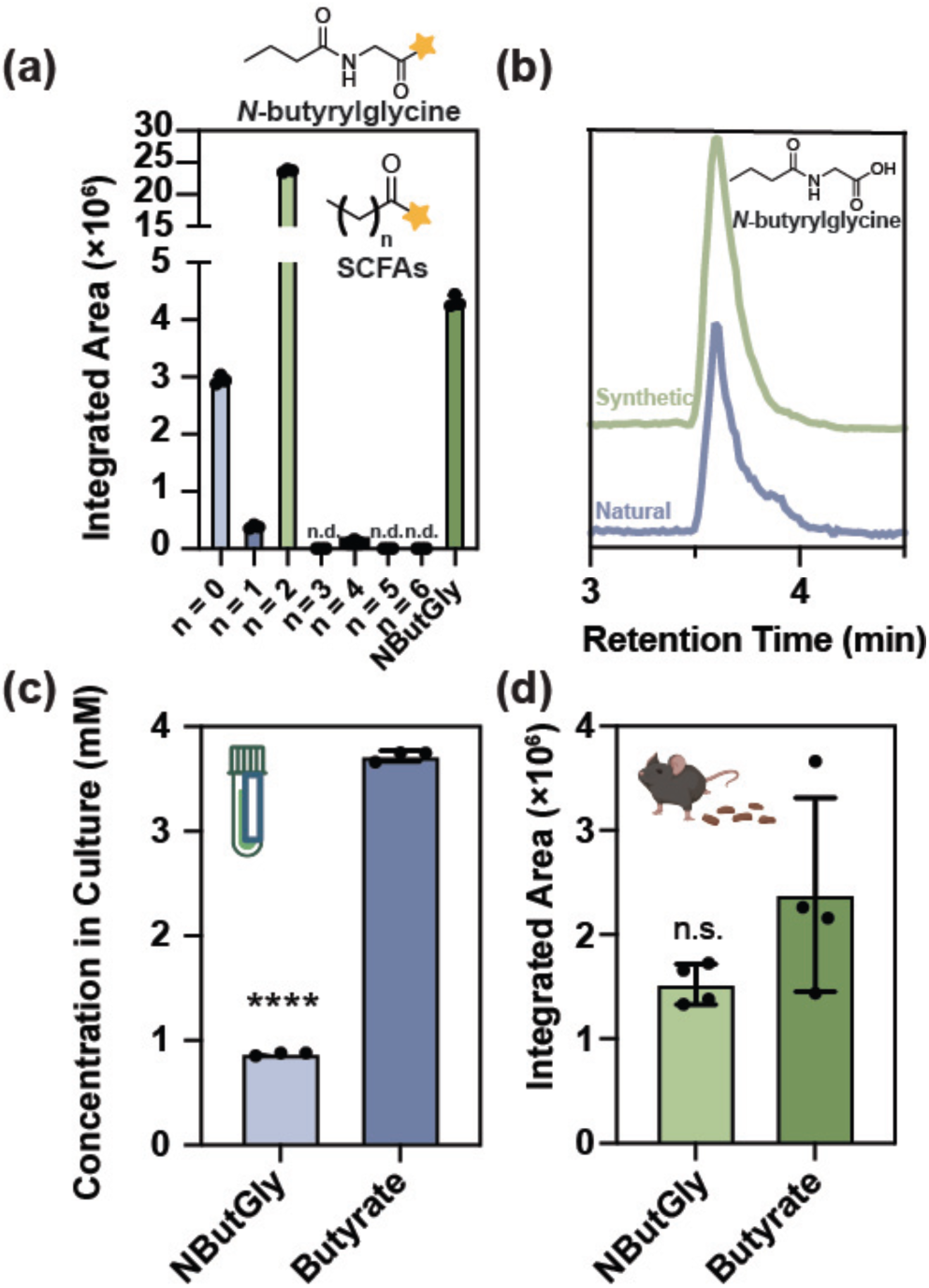
*Faecalibacterium prausnitzii* produces the metabolite *N*-butyrylglycine. (a) LC-MS analysis of 3-NPH-derivatized short-chain fatty acids (SCFAs) and *N*-butyrylglycine produced by *F. prausnitzii*. Chemical structures are shown. SCFAs with chain lengths of n = 0, 1, 2, 3, 4, 5, and 6 correspond to 3-NPH analogs of acetate, propionate, butyrate, pentanoate, hexanoate, heptanoate, and octanoate, respectively. n.d. stands for not detected. (b) Representative traces for the comparison of synthetic and naturally occurring *N*-butyrylglycine by LC-MS retention time analysis. (c) LC-MS quantification of *N*-butyrylglycine and butyrate in stationary-phase *F. prausnitzii* cultures. (d) LC-MS detection of *N*-butyrylglycine and butyrate in fecal samples from conventional wildtype mice (n = 4 biological replicates). Data are presented as mean ± SD. Statistical significance was determined using Welch’s t-test; *P* ≤ 0.0001 (****), n.s., not significant. All data represent at least three biological replicates unless otherwise stated.

*N*-butyrylglycine has been reported as an endogenous mammalian metabolite and biomarker of altered fatty acid metabolism,^20–21^ yet little is known regarding its biological activity. It belongs to the broader family of *N*-acyl amides, a privileged class of signaling metabolites that have emerged as important regulators of host physiology.^22,23^ The identification of NButGly in *F. prausnitzii* cultures therefore prompted us to investigate the levels at which it is produced. Quantitative LC-MS analysis showed that NButGly accumulated to concentrations approaching 1 mM in stationary-phase cultures, while butyrate reached concentrations of roughly 3.7 mM (**Fig. 2c**). Moreover, to determine whether NButGly is present within the mammalian gut environment, we analyzed fecal samples from conventional wildtype mice. Both NButGly and butyrate were readily detected in these samples (**Fig. 2d**), indicating that NButGly is a naturally occurring component of the gut microbiome. Notably, while NButGly has traditionally been regarded as a mammalian metabolite,^20,21^ our findings identify *F. prausnitzii* as a microbial source of this molecule and suggest that the gut microbiota contributes to its production, although the microbial origins of NButGly detected in murine feces remain undetermined.

In humans, *N*-acylglycines can be synthesized by the mitochondrial enzyme glycine *N*-acyltransferase (GLYAT), which catalyzes the conjugation of glycine with acyl-CoA substrates, including butyryl-CoA.^24^ However, a BLASTp search using human GLYAT against the *F. prausnitzii* genome did not identify a detectable homolog, suggesting that *N*-butyrylglycine production in *F. prausnitzii* may involve a distinct *N*-acyltransferase. Given that NButGly is an *N*-acyl amide harboring a butyryl moiety, we hypothesized that its biosynthesis is linked to butyrate production in *F. prausnitzii*. In this bacterium, butyrate is synthesized through a reverse β-oxidation pathway, in which acetate is consumed to produce butyryl-CoA before conversion to butyrate (**Fig. 3a**).^25^ Since butyryl-CoA is both a central intermediate in butyrate biosynthesis and a high-energy thioester, we reasoned that NButGly could be generated by *N*-acyltransferase-catalyzed condensation of butyryl-CoA and glycine. To identify candidate enzymes capable of catalyzing this reaction, we performed an in silico survey of putative *N*-acyltransferases encoded by the *F. prausnitzii* genome. Reasoning that a candidate enzyme may preferentially bind butyryl-CoA, we docked^26^ both butyryl-CoA and acetyl-CoA into predicted structures of 20 candidate *N*-acyltransferases and prioritized candidates based on their relative binding preference for the two acyl donors (**Table S1**). Heterologous expression and in vitro biochemical characterization of selected candidates identified ACOP_v1_740045 (**Fig. S6**) as an *N*-butyryltransferase capable of catalyzing NButGly formation in vitro (**Fig. 3b**). Enzymatic reactions were performed using 20 µM purified enzyme, 0.5 mM butyryl-CoA, and 10 mM glycine for 1 h at room temperature. In the absence of enzyme, low levels of NButGly were detected, consistent with spontaneous non-enzymatic acyl transfer from butyryl-CoA to glycine. Under these conditions, addition of ACOP_v1_740045 increased NButGly production by approximately 9-fold relative to the enzyme-free control. These findings establish the biochemical capacity of ACOP_v1_740045 to catalyze NButGly formation, although its contribution to NButGly biosynthesis in *F. prausnitzii* remains to be determined.

**Fig. 3.**
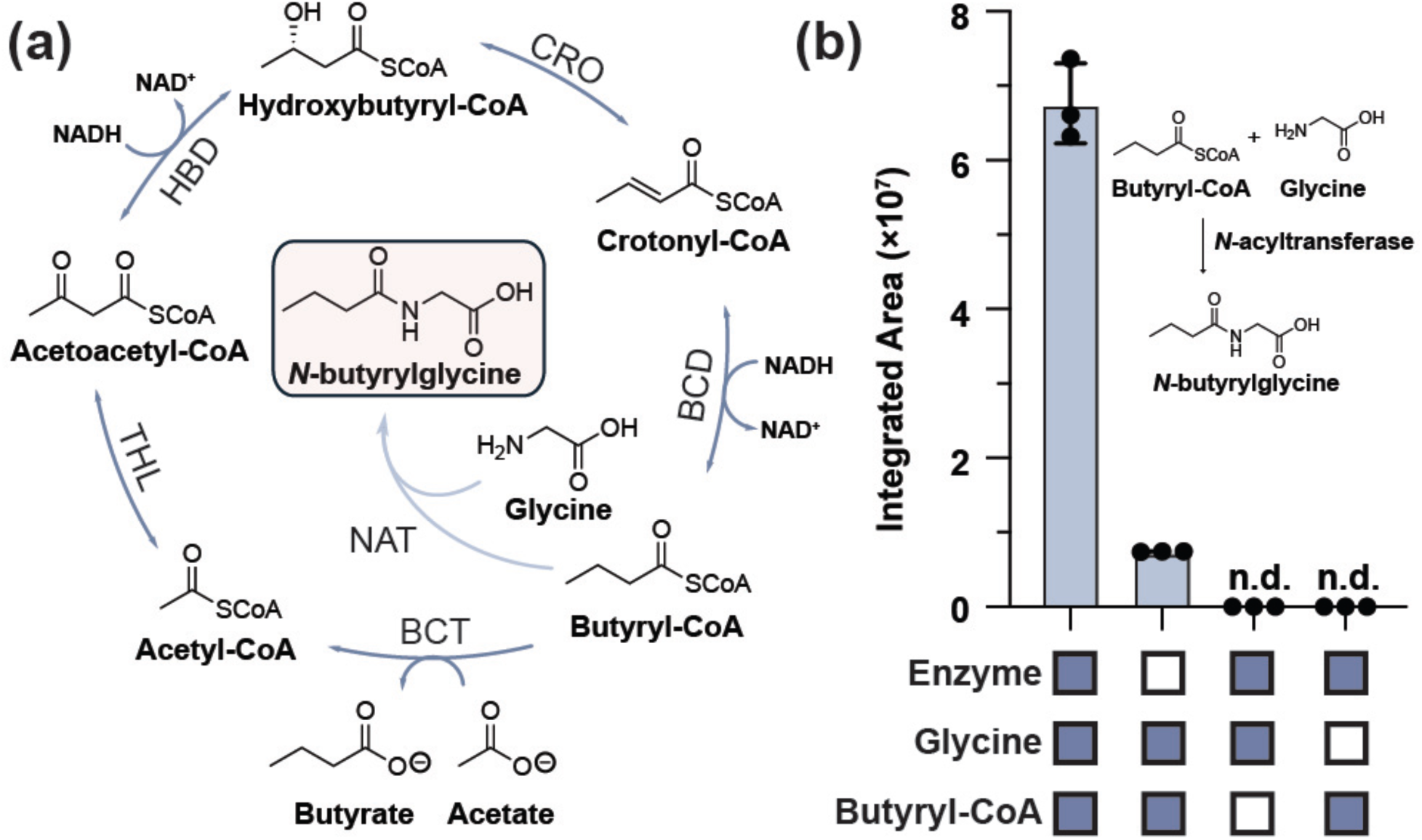
Identification of an *N*-acyltransferase that catalyzes *N*-butyrylglycine biosynthesis. (a) Proposed biosynthetic route linking butyrate metabolism to *N*-butyrylglycine production in *F. prausnitzii*. THL, HBD, CRO, BCD, BCT, and NAT denote thiolase, 3-hydroxybutyrate dehydrogenase, crotonase, butyryl-CoA dehydrogenase, butyryl-CoA:acetate CoA-transferase, and *N*-acyltransferase, respectively. (b) In vitro reconstitution of *N*-butyrylglycine biosynthesis using purified ACOP_v1_740045, glycine, and butyryl-CoA. Blue and white boxes indicate the presence and absence, respectively, of each reaction component. Data are presented as mean ± SD; n.d., not detected, n = 3 technical replicates.

Next, we examined whether ACOP_v1_740045 could utilize acetyl-CoA as an acyl donor and accept other amino acid substrates. Using acetyl-CoA or butyryl-CoA, we evaluated the acylation of glycine, alanine, valine, leucine, isoleucine, proline, and phenylalanine. We found that ACOP_v1_740045 catalyzed both acetylation and butyrylation of glycine, whereas no detectable activity was observed with the alternative amino acids. Steady-state kinetic analyses revealed apparent Kₘ values for glycine of 61.95 and 485 µM with acetyl-CoA and butyryl-CoA, respectively (**Fig. S7**). Although butyrylation exhibited a higher apparent turnover number (k_cat_, 1.88 versus 1.05 s⁻¹), acetylation displayed approximately 4.3-fold greater apparent catalytic efficiency (k_cat_/K_m_) with respect to glycine. Together, these findings demonstrate that ACOP_v1_740045 selectively acylates glycine among the examined amino acids, with distinct kinetic preferences for acetyl-CoA and butyryl-CoA.

To investigate whether homologs of ACOP_v1_740045 are present in other human-associated bacteria, we performed a BLASTp search on the Human Microbiome Project (HMP) reference genomes.^27^ This analysis identified eight homologs, including closely related sequences in *Faecalibacterium* and more divergent sequences in *Subdoligranulum* and *Anaerotruncus* (**Table S2**). An expanded search of the MGnify Unified Human Gastrointestinal Protein^28^ Catalog identified 78 significant protein matches across diverse gut-associated bacterial taxa, including 32 sequences from *Faecalibacterium*, 27 from *Gemmiger*, as well as additional homologs in *Ruthenibacterium*, *Fournierella*, and other genera (**Table S3**). We also searched the MGnify Mouse Gut Protein Catalogue for homologs in the murine gut microbiome. This analysis identified 28 significant homologs across several bacterial genera, including seven highly conserved sequences (64-97% amino acid identity) from *Faecalibacterium* species (**Table S4**). These findings demonstrate that homologs of ACOP_v1_740045 are not restricted to *Faecalibacterium*.

Having established both the enzymatic production and millimolar accumulation of NButGly in *F. prausnitzii* cultures, we next sought to determine how glycine conjugation to butyrate influences its bioactivity. Butyrate is a principal metabolite of the gut microbiome that regulates host physiology through diverse mechanisms, including weak, reversible inhibition of histone deacetylase (HDACs).^29^ We therefore began by comparing HDAC inhibitory activities of butyrate and NButGly. Consistent with its butyrate-derived structure, NButGly inhibited HDAC activity in a dose-dependent manner, although it was less potent than butyrate across all tested concentrations (**Fig. 4a**). Nonlinear regression analysis yielded estimated IC_50_ values of 2.824 mM for NButGly and 126.8 µM for butyrate, indicating an approximately 22-fold difference in inhibitory potency. These findings suggest that glycine conjugation attenuates the canonical biological activity of butyrate by diminishing its ability to inhibit HDACs.

**Fig. 4.**
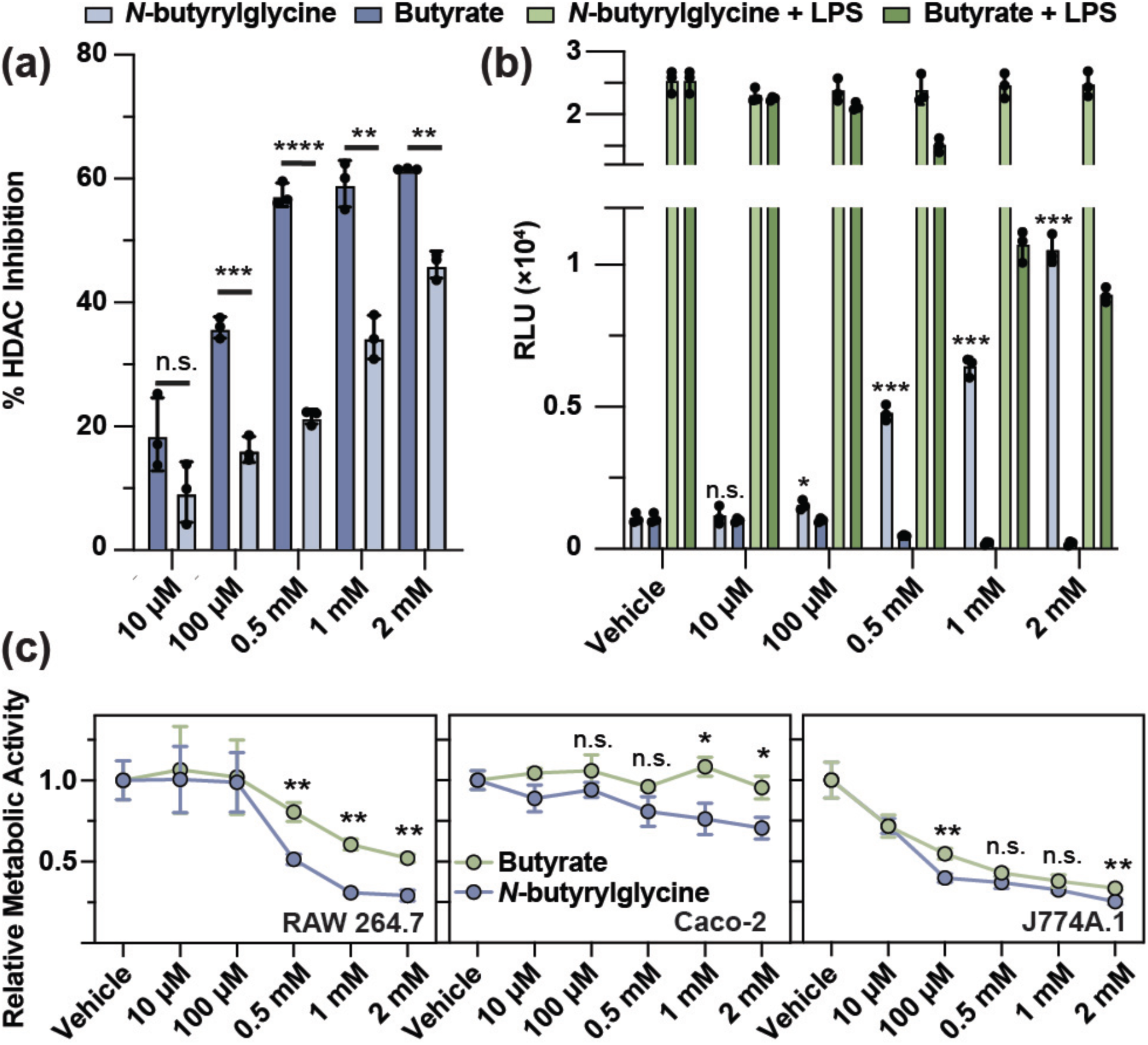
*N*-butyrylglycine exhibits biological activities distinct from butyrate. (a) Comparison of histone deacetylase (HDAC) inhibitory activity of *N*-butyrylglycine and butyrate across the indicated concentrations. (b) Effects of *N*-butyrylglycine and butyrate on NF-κB signaling in LPS-stimulated and unstimulated RAW 264.7 macrophages using an NF-κB luciferase reporter cell line. Statistical comparisons were performed between each unstimulated *N*-butyrylglycine concentration and the corresponding vehicle control. (c) Dose-dependent effects of *N*-butyrylglycine and butyrate on metabolic activity in RAW 264.7 macrophages, Caco-2 colorectal adenocarcinoma cells, and J774A.1 macrophages. Relative metabolic activity was determined by the 3-(4,5-dimethylthiazol-2-yl)-2,5-diphenyltetrazolium bromide (MTT) assay. Data are presented as mean ± SD. Statistical significance was determined using Welch’s t-test; *P* ≤ 0.05 (*), *P* ≤ 0.01 (**), *P* ≤ 0.001 (***), *P* ≤ 0.0001 (****), n.s., not significant. All data represent at least three biological replicates.

Since HDAC inhibition is closely linked to the anti-inflammatory properties of butyrate,^30^ we next examined the effects of NButGly on NF-κB signaling using a RAW 264.7 reporter cell line.^31^ Consistent with previous reports, butyrate suppressed lipopolysaccharide (LPS)-induced NF-κB activation in a concentration-dependent manner (**Fig. 4b**).^32^ In contrast, NButGly failed to attenuate LPS-induced signaling and, surprisingly, induced NF-κB reporter activity in unstimulated macrophages at concentrations above 0.5 mM. Together with its weaker HDAC inhibitory activity, these findings indicate that glycine conjugation fundamentally alters the biological activities of butyrate.

Given the established role of butyrate in regulating cellular proliferation,^33,34^ we next investigated whether NButGly similarly influences mammalian cell growth. Using an MTT assay, we found that treatment with NButGly resulted in a dose-dependent reduction in metabolic activity across select mammalian cell lines, including RAW 264.7 macrophages, J774A.1 macrophages, and Caco-2 colorectal adenocarcinoma cells (**Fig. 4c**). We found that NButGly reduced metabolic activity at lower concentrations and with greater potency than butyrate in these cell lines, despite exhibiting weaker HDAC inhibition and distinct effects on inflammatory signaling. Importantly, while both NButGly and butyrate inhibited cell proliferation, trypan blue exclusion assays confirmed that neither compound significantly reduced cell viability at 2 mM relative to untreated controls (**Fig. S8**). Collectively, these findings demonstrate that NButGly is a bioactive metabolite with biological activities distinct from its parent SCFA.

Since depletion of *F. prausnitzii* has been associated with CRC incidence,^6,7^ we were interested in deciphering the molecular basis underlying the anti-proliferative phenotype of NButGly. To this end, we performed RNA-seq transcriptional profiling on Caco-2 colorectal adenocarcinoma cells treated with either 0.5 mM NButGly or vehicle control. Gene set enrichment analysis (GSEA) of this dataset revealed significant downregulation of the G2/M checkpoint and E2F target pathways (**Fig. 5a**, **Table S5**). The G2/M checkpoint and E2F transcriptional programs coordinate cell-cycle progression, DNA replication, and mitotic entry, and their dysregulation is a defining feature of many human cancers.^35,36^ Their coordinated repression therefore suggests that NButGly suppresses proliferative signaling by limiting progression through the cell cycle. Consistent with this model, qRT-PCR confirmed significant downregulation of *MKI67*, a canonical marker of actively proliferating cells and a downstream target of cell-cycle regulatory pathways (**Fig. 5b**).^37^ Importantly, butyrate treatment similarly downregulated *MKI67*, in agreement with its established anti-proliferative activity (**Fig. S9**).^38^ To further investigate the effects of NButGly on cell-cycle progression, we analyzed cellular DNA content using propidium iodide staining and flow cytometry. NButGly treatment increased the relative prominence of the 4N DNA peak compared with the 2N peak (**Fig. S10**), consistent with accumulation of cells in G2/M. These findings suggest that the altered DNA-content distribution may reflect impaired progression through G2/M, in line with the observed downregulation of cell-cycle-associated transcriptional programs. Together, these findings suggest that the anti-proliferative activity of NButGly is accompanied by repression of cell-cycle-associated transcriptional programs.

**Fig. 5.**
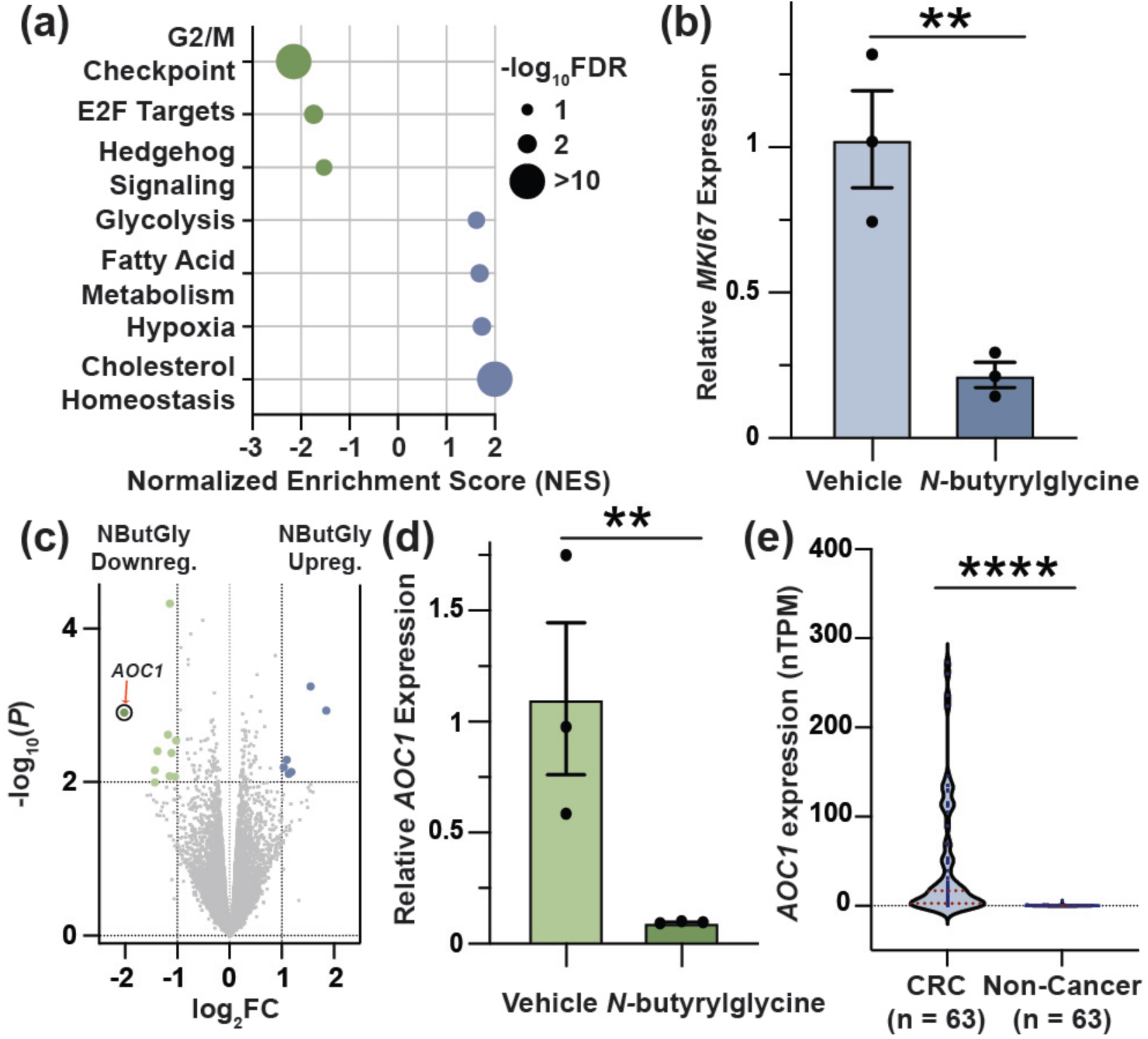
*N*-butyrylglycine represses colorectal cancer-associated transcriptional programs. (a) Gene set enrichment analysis (GSEA) of RNA-seq data from Caco-2 colorectal adenocarcinoma cells treated with 0.5 mM *N*-butyrylglycine or vehicle control. Bubble size corresponds to pathway significance (-log_10_FDR), while bubble color indicates pathways downregulated (green; negative NES) or upregulated (blue; positive NES) following *N*-butyrylglycine treatment. (b) qRT-PCR validation of the proliferation marker *MKI67* following *N*-butyrylglycine treatment. (c) Volcano plot analysis of genes differentially regulated following *N*-butyrylglycine treatment. Differentially expressed genes meeting significance thresholds (-log_10_(*P*) ≥ 2, |log_2_(FC)| ≥ 2) are colored green (downregulated by NButGly) or blue (upregulated by NButGly); *AOC1* is circled and indicated with an arrow. Fold change (FC) was calculated as the ratio of transcript abundance in vehicle-treated cells relative to NButGly-treated cells. (d) qRT-PCR validation of *AOC1* downregulation following *N*-butyrylglycine treatment. (e) *AOC1* expression in colorectal cancer and non-cancerous cell lines obtained from publicly available transcriptomic datasets. Measurements for individual cell lines are shown as blue points, while red dashed lines indicate the median and interquartile range. Data are presented as mean ± SEM. Statistical significance was determined using Welch’s t-test; *P* ≤ 0.01 (**), *P* ≤ 0.0001 (****). All data represent at least three biological replicates.

In addition to pathway-level changes, differential gene expression analysis identified several genes that were strongly regulated by NButGly treatment (**Fig. 5c**, **Table S6**). Interestingly, NButGly elicited a relatively limited transcriptional response compared with the broad changes previously reported following butyrate treatment in Caco-2 cells. Studies using 3 mM butyrate have identified thousands of differentially expressed genes,^39^ whereas treatment with 0.5 mM NButGly resulted in substantially fewer significant transcriptional changes. This difference may reflect the lower treatment concentration used in our study, as well as the weaker HDAC inhibitory activity of NButGly relative to butyrate. However, the possibility that NButGly influences additional regulatory pathways independent of HDAC inhibition cannot be excluded.

Among the genes regulated by NButGly, amine oxidase copper-containing 1 (*AOC1*) was the most robustly downregulated transcript, a finding independently validated by qRT-PCR (**Fig. 5d)**. Notably, butyrate treatment did not significantly alter *AOC1* levels relative to control (**Fig. S11**), suggesting that NButGly may elicit transcriptional responses distinct from those of butyrate. AOC1 is a copper-dependent amine oxidase that contributes to the degradation of polyamines and histamine through oxidative deamination and has emerged as an oncogenic factor in human cancers.^40,41^ In CRC, elevated *AOC1* expression has been associated with tumor progression, and genetic knockdown of *AOC1* has been shown to suppress CRC cell proliferation and tumor growth.^41,42^ Consistent with these observations, analysis of publicly available transcriptomic datasets show that *AOC1* expression is markedly elevated in CRC cell lines relative to non-cancerous cell lines (**Fig. 5e**, **Table S7**).^43^ Together, these findings identify *AOC1* as a transcriptional target of NButGly and further suggest that microbial production of this compound may contribute to the established relationship between *F. prausnitzii* intestinal abundance and CRC.

Collectively, our findings establish *N*-butyrylglycine as a robustly produced metabolite of *F. prausnitzii* that is readily detectable in fecal samples from conventional wildtype mice and exhibits biological activities that differ from those of its parent SCFA, butyrate. In addition to identifying the biosynthetic enzyme responsible for NButGly production, our data demonstrate that this metabolite suppresses proliferative transcriptional programs and downregulates the CRC-associated gene *AOC1*. Together, these findings reveal that *F. prausnitzii* not only produces butyrate as a major bioactive metabolite, but also enzymatically remodels it into a chemically and biologically distinct molecule. More broadly, our work suggests that deciphering the secondary metabolism of *F. prausnitzii* may be critical for understanding the molecular basis through which this health-associated bacterium influences CRC and other disorders linked to intestinal dysbiosis.

## Data Availability Statement

The datasets supporting I this article have been uploaded as part of the supporting information: Complete Materials and Methods, NMR, HR-MS, and HR-MS/MS spectra, and associated references (PDF).

AutoDock Vina binding energy calculations (Table S1), ACOP_v1_740045 BLASTp results against HMP reference genomes (Table S2), ACOP_v1_740045 BLASTp results against MGnify Human Gut Protein Catalogue (Table S3), ACOP_v1_740045 BLASTp results against MGnify Mouse Gut Protein Catalogue (Table S4), Gene Set Enrichment Analysis (Table S5), differential gene expression analysis (Table S6), and *AOC1* transcript abundances across cell lines (Table S7) (XLSX).

## ACKNOWLEDGMENT

This work was supported by an NIH R35 Maximizing Investigators’ Research Award for Early Stage Investigators (R35GM133501 to P.V.C.), a Beckman Young Investigator Award from the Arnold and Mabel Beckman Foundation (to P.V.C.), a Sloan Research Fellowship from the Alfred P. Sloan Foundation (to P.V.C.), and a National Science Foundation Postdoctoral Research Fellowship in Biology (Award No. 2507806 to R.H.). This work made use of the Cornell University NMR Facility, which is supported, in part, by the NSF through MRI award CHE-1531632. We are grateful to the Weill Institute for Cell and Molecular Biology for additional resources. Portions of figures in this manuscript were generated with BioRender (Agreement number: JT29XF1F3I).^44^

