## Supplementary Information for "Gut Microbial Butyrate Metabolism Generates an *N*-Acyl Amide that Regulates Host Transcriptional Programs"

#### Table of Contents

**Methods** 3
**General Experimental Procedure** 3
***F. prausnitzii* Culture and Extraction** 3
**Extraction of Metabolites from Mouse Fecal Samples** 3
**3-NPH Derivatization of Bacterial and Fecal Samples** 4
**LC-MS Analysis of Bacterial and Fecal Samples** 4
**Synthesis of *N*-butyrylglycine** 4
**Histone Deacetylase (HDAC) Inhibition Assay** 4
**NF-κB Luciferase Activity Assay** 5
**MTT Proliferation Assay** 5
**Trypan Blue Exclusion Assay** 5
**RNA-Seq Sample Preparation and Analysis** 6
**Quantitative RT-PCR of Mammalian Genes** 6
**Protein Structure Prediction and Molecular Docking** 6
**Purification of ACOP_v1_740045** 7
**In vitro Enzymatic Assays for *N*-butyrylglycine Biosynthesis** 7
**Steady-State Kinetic Analysis of ACOP_v1_740045** 7
**Cell-Cycle Analysis by Propidium Iodide Staining and Flow Cytometry** 8
**Figure S1**. ESI$^-$-HRMS/MS fragmentation spectrum of natural 3-NPH-derivatized *N*-butyrylglycine (20 eV). Structures consistent with fragmentation pattern are shown. 10
**Figure S2**. Synthesis of *N*-butyrylglycine. 11
**Figure S3**. $^1$H NMR spectrum of *N*-butyrylglycine (400 MHz, methanol-$d_4$). 12
**Figure S4**. $^{13}$C NMR spectrum of *N*-butyrylglycine (400 MHz, methanol-$d_4$). 13
**Figure S5**. ESI$^-$-HRMS/MS fragmentation spectrum of synthetic (top) and natural (bottom) 3-NPH-derivatized *N*-butyrylglycine (20 eV). Structures consistent with fragmentation pattern are shown. 14
**Figure S6**. Purification of ACOP_v1_740045. 15
**Figure S7**. Steady-state kinetic characterization of ACOP_v1_740045-mediated glycine acylation. 16
**Figure S8**. Effects of NButGly and butyrate treatment on mammalian cell viability. 17
**Figure S9**. Effect of butyrate treatment on *MKI67* expression in Caco-2 cells. 18
**Figure S10**. Effect of NButGly treatment on cellular DNA-content distribution in Caco-2 cells. 19
**Figure S11**. Effect of butyrate treatment on *AOC1* expression in Caco-2 cells. 20
**References** 21

### Methods

#### General Experimental Procedure

All reagents used were analytical grade and obtained from commercial suppliers without further purification. High-resolution electrospray ionization mass spectrometry (HR-ESI-MS) was performed on an Agilent 6230 electrospray ionization-time-of-flight (ESI-TOF) mass spectrometer coupled to an Agilent 1290 Infinity HPLC system equipped with an InfinityLab Poroshell 120 EC-C18 column (3.0 × 50 mm, 2.7 µm). Chromatography was performed using a linear gradient of water and acetonitrile, both containing 0.1% formic acid at a flow rate of 0.5 mL/min. The gradient increased from 10% to 100% acetonitrile over 10 min, followed by a 4 min 100% acetonitrile wash. Mass spectra were acquired in negative ionization mode over an *m/z* range of 100-1700. Targeted MS/MS analyses were performed using an isolation width of 1.3 *m/z* and a fixed collision energy of 20 eV. NMR spectra were obtained on a 400 MHz Bruker ACIII HD spectrometer equipped with a broadband Fluorine Observe Probe at ambient temperature. Mammalian cells were cultured in Dulbecco's Modified Eagle Medium (DMEM) supplemented with 10% fetal bovine serum (FBS) and 1% penicillin-streptomycin at 37 ˚C in a 5% $CO_2$ atmosphere. Fluorescence, luminescence, and absorbance measurements were acquired using a BioTek Synergy H1 microplate reader. Flow cytometry measurements were acquired using a BD Accuri C6 Plus flow cytometer (BD Biosciences), and data were analyzed using BD Accuri software. All experiments were performed at least three independent times. RAW 264.7, Caco-2, and J774A.1 cells were purchased from the ATCC repository. RAW 264.7 cells stably expressing an NF- kB responsive firefly luciferase reporter[1] were a kind gift from the Medzhitov lab at Yale University.

#### *F. prausnitzii* Culture and Extraction

A starter culture of *F. prausnitzii* was grown anaerobically in 5 mL of Reinforced Clostridial Medium (RCM) at 37˚C without agitation for 48 h. The culture was diluted 1:100 into fresh RCM (5 mL) in biological triplicate and incubated under identical conditions for an additional 48 h. Cultures were centrifuged at 4,500 × g for 10 min to separate the bacterial cells from the culture supernatant. Culture supernatants were collected directly and subjected to 3-nitrophenylhydrazine (3-NPH) derivatization. Cell pellets were resuspended in 2 mL of methanol:acetonitrile:water (2:2:1, v/v/v), sonicated for 5 min, and incubated at room temperature for 1 h. Cell debris was removed by centrifugation (17,000 × g, 10 min), and the resulting supernatants were collected for 3-NPH derivatization.

#### Extraction of Metabolites from Mouse Fecal Samples

Animal studies were approved by and performed in accordance with the guidelines of the Institutional Animal Care and Use Committee (Protocol number 2015-0069). Mouse fecal pellets were collected from conventional wildtype M/F C57Bl6 mice, age 8-10 weeks (Jackson Labs) and frozen until further use. Fecal samples (1 g) were resuspended in 5 mL of methanol:acetonitrile:water (2:2:1, v/v/v), sonicated for 10 min, and incubated at 37˚C with shaking (250 rpm) for 1 h. Samples were centrifuged at 4,500 × g, 10 min to pellet insoluble material, and the resulting supernatants were collected for 3-NPH derivatization.

##### 3-NPH Derivatization of Bacterial and Fecal Samples

For metabolomic analysis, 10 µL of bacterial pellet extract and 10 µL of bacterial supernatant was mixed with 10 µL of 200 mM 3-NPH in 50% aqueous acetonitrile and 10 µL of 120 mM 1-ethyl-3-(3-dimethylaminopropyl)carbodiimide (EDC) prepared in 50% aqueous acetonitrile containing 6% pyridine, to form a 40 µL solution. The reaction mixture was incubated at 42˚C for 30 min and diluted to a final volume of 200 µL with 10% aqueous acetonitrile. Samples were clarified by centrifugation (17,000 × g, 10 min), and 1 µL of the resulting supernatant was analyzed by LC-MS. For mouse fecal extracts, 20 µL of extract was subjected to derivatization in place of bacterial samples.

##### LC-MS Analysis of Bacterial and Fecal Samples

Metabolite concentrations in bacterial cultures were determined using calibration curves generated from synthetic *N*-butyrylglycine standards that were derivatized in parallel under identical conditions. Extracted ion chromatograms (EICs) were generated using the exact monoisotopic deprotonated mass ($[M – H]^-$) with a mass tolerance of 10 ppm. Peak integration was performed using Agilent MassHunter Qualitative Analysis software.

##### Synthesis of *N*-butyrylglycine

Glycine (180 mg, 2.4 mmol) was dissolved in 10 mL of 1 M NaOH and cooled on ice for 30 min. Butyryl chloride (270 µL, 2.6 mmol) was added dropwise, and the reaction mixture was allowed to warm to room temperature and stirred for 16 h. The reaction was acidified to pH 2 with HCl and extracted with ethyl acetate (3 × 20 mL). The combined organic extracts were concentrated via rotary evaporation and dried under high vacuum using a Schlenk line to remove residual butyric acid affording *N*-butyrylglycine as a white film (35 mg, 10%).

*N*-butyrylglycine: $C_6H_{11}NO_3$; white film; $^1$H NMR (400 MHz, MeOD-$d_4$) δ 3.82 (s, 2H), 2.13 (t, $J$ = 7.63 Hz, 2H), 1.60 (m, 2H), 0.90 (t, $J$ = 7.48, 3H); $^{13}$C NMR (100 MHz, MeOD-$d_4$) δ 175.17, 171.69, 40.33, 37.30, 18.81, 12.55.; LC-MS (ESI, negative mode) of the 3-NPH derivative: calcd. *m/z* 279.1099 ($C_{12}H_{15}N_4O_4^-$, $[M – H]^-$); found: *m/z* 279.1095.

##### Histone Deacetylase (HDAC) Inhibition Assay

Caco-2 cells were cultured to 80-90% confluence in 10 cm tissue culture dishes, detached with 0.05% trypsin for 3 min at 37˚C, and collected by centrifugation (350 × g, 3 min, room temperature). Cell pellets were resuspended in Nonidet P-40 lysis buffer (25 mM Tris-HCl, pH 8.0, 150 mM NaCl, 10% glycerol, and 1% Nonidet P-40) supplemented with protease inhibitor cocktail (1:100 dilution) and sonicated to ensure complete lysis. Cell debris was removed by centrifugation (17,000 × g, 10 min, 4˚C), and the resulting whole-cell lysates were collected. Protein concentrations were determined using the Bio-Rad DC Protein Assay.

HDAC activity assays were performed in 55 μL reaction volumes. Whole-cell lysates (normalized to 1 mg/mL by DC Assay) were first incubated with *N*-butyrylglycine, butyrate, or solvent vehicle control (10% aqueous DMSO containing 5% methanol) for 30 min at 37˚C in reaction buffer (25 mM Tris-HCl, pH 8.0, 50 mM NaCl, and 1 mM DTT). Following pre-incubation, the fluorogenic HDAC substrate BOC-Lys(Ac)-AMC was added to a final concentration of 100 μM, and the reactions were incubated for an additional 1 h at 37˚C. Trypsin solution (55 μL, 10 mg/mL) was then added, and the reactions were incubated for 20 min at 37˚C. Aliquots (100 μL) were transferred to black 96-well plates, and fluorescence was measured using excitation and emission wavelengths of 355 and 460 nm, respectively.

**NF-κB Luciferase Activity Assay**

RAW 264.7 macrophages stably expressing an NF-kB responsive firefly luciferase reporter[1] were seeded at $10^5$ cells/well in a 24-well plate. After 48 h (70-80% confluence) the culture medium was replaced, and cells were treated with *N*-butyrylglycine, butyrate, or vehicle control (10% aqueous DMSO containing 5% methanol) for 3 h at 37˚C and 5% $CO_2$. Cells were washed once with phosphate-buffer saline (PBS) and lysed with 1X Passive Lysis Buffer (25 mM Tris-HCl, pH 7.8, 2 mM 1,2-diaminocyclohexane tetraacetic acid, 2 mM DTT, 1 mg/mL BSA, 1% (v/v) Triton X-100, and 10% (v/v) glycerol in dd$H_2O$). Aliquots (20 μL) of cell lysate were transferred to white opaque 96-well plates and mixed with 50 μL of firefly luciferase substrate solution (75 mM HEPES, pH 8.0, 4 mM $MgSO_4$, 20 mM DTT, 0.1 mM EDTA, 0.5 mM ATP, 0.25 mM coenzyme A, and 0.5 mM D-luciferin (firefly) in dd$H_2O$). Luminescence was measured immediately using a plate reader.

For LPS stimulation experiments, cells were cultured as described above and pre-incubated with *N*-butyrylglycine, butyrate, or vehicle control for 1 h prior to addition of LPS (100 ng/mL) or solvent control. Cells were incubated an additional 3 h at 37˚C and 5% $CO_2$, after which luciferase activity was quantified as described above.

**MTT Proliferation Assay**

Cells were seeded at $5\times10^4$ cells/well in 96-well tissue culture plates. After 24 h, the cells were treated with *N*-butyrylglycine, butyrate, or vehicle control (10% aqueous DMSO containing 5% methanol) for 24 h at 37˚C and 5% $CO_2$. Afterwards, the culture medium was replaced with fresh medium containing 1.2 mM MTT, and plates were incubated for an additional 3 h. Cell-free wells containing MTT reagent were included as background controls. Following incubation, 75 μL of medium was removed from each well and replaced with 50 μL of DMSO to dissolve the resulting formazan crystals (10 min, 37˚C). Aliquots (50 μL) were transferred to clear-bottom 96-well plates, and UV absorbance was measured at 540 nm using a plate reader.

**Trypan Blue Exclusion Assay**

Cells were seeded at $5 \times 10^4$ cells/well in 96-well tissue culture plates. After 24 h, cells were treated with 2 mM *N*-butyrylglycine, 2 mM butyrate, or vehicle control (0.1% aqueous DMSO containing 0.05% methanol) for 24 h at 37 °C and 5% $CO_2$. Following treatment, culture medium was removed, and cells were washed once with PBS before detachment with 0.05% trypsin (3 min, 37 °C). Trypsinization was quenched with fresh culture medium, and cells were collected by centrifugation (350 × g, 3 min). Cells were subsequently resuspended and stained with trypan blue solution. Viable and nonviable cells were counted using a hemocytometer, and cell viability was calculated as the percentage of viable cells relative to the total number of cells counted. Viability measurements were normalized to the vehicle-treated control.

**RNA-Seq Sample Preparation and Analysis**

Caco-2 cells were seeded at 1 × $10^5$ cells/well in 6-well tissue culture plates and cultured for 30 h at 37˚C and 5% $CO_2$. Cells were treated with either 0.5 mM *N*-butyrylglycine or vehicle control (10% aqueous DMSO containing 5% methanol) for 6 h (n = 3 biological replicates per condition). Following treatment, culture medium was removed, and cells were washed once with PBS before detachment with 0.05% trypsin (3 min, 37˚C). Trypsinization was quenched with fresh culture medium, and cells were collected by centrifugation (350 × g, 3 min). Cell pellets were resuspended in 100 µL of 1X DNA/RNA Shield (Zymo Research), vortexed for 2 min, and submitted to Plasmidsaurus (Eugene, OR, USA) for RNA-seq and transcriptomic analysis. Library preparation, sequencing, read alignment, differential gene expression analysis, and gene set enrichment analysis were performed using the Plasmidsaurus standard RNA sequencing workflow according to the manufacturer's protocols. Detailed information regarding, sequencing, quality control, and bioinformatic analysis is available in the Plasmidsaurus RNA-seq technical documentation (https://plasmidsaurus.com/technical-documentation/rna).

**Quantitative RT-PCR of Mammalian Genes**

Caco-2 cells were cultured and treated with 0.5 mM *N*-butyrylglycine, butyrate, or vehicle control as described for RNA-sequencing. Total RNA was isolated using RNA Bee (Tel-Test) according to the manufacturer's protocol with glycogen added during isopropanol precipitation to improve RNA recovery. RNA concentrations were determined by NanoDrop spectrophotometry, and 1 µg of total RNA was reverse transcribed using Moloney murine leukemia virus reverse transcriptase (MMLV RT, Takara Bio) and oligo(dT) primers. Quantitative PCR was performed using QuantaBio SYBR Green master mix with gene-specific primers on a Bio-Rad Real-Time PCR Detection System. Relative transcript abundance was calculated using the ΔΔCt method following normalization to the housekeeping gene *RPLP0.*

**Protein Structure Prediction and Molecular Docking**

Protein structures were predicted from amino acid sequences using ColabFold.[2] For each protein, the top-ranked predicted structure was selected for docking analyses using the relaxed model. Predicted protein structures were converted from PDB to PDBQT format using Open Babel.[3] Polar hydrogens were added during receptor preparation, and Gasteiger partial charges were assigned to all receptor atoms. Ligands were generated from SMILES strings, converted to three-dimensional structures using Open Babel, and subsequently converted to PDBQT format using Meeko.[4] Molecular docking was performed using AutoDock Vina v1.2.7.[5] An initial blind docking calculation was performed using adenosine diphosphate (ADP) as a probe ligand and a search box encompassing the entire predicted protein structure to identify a putative nucleotide-binding site. The centroid of the highest-scoring docked pose was subsequently used to define a focused docking box (24 × 24 × 24 Å) for all subsequent docking calculations. Truncated acetyl-CoA and butyryl-CoA analogs containing only the acyl-thioester and pantetheine arm were docked into the same focused search space for each protein to enable direct comparison. Docking scores (kcal/mol) reported by the AutoDock Vina scoring function were used to compare the relative binding preferences of candidate *N*-acyltransferases.

**Purification of ACOP_v1_740045**

The *F. prausnitzii ACOP_v1_740045* gene was codon optimized for expression in *E. coli* and ordered from Twist Bioscience. The gene was cloned into a pET-28a vector using XhoI and EcoRI, resulting in a C-terminal $His_{6x}$-tag fusion, and transformed into *E. coli* BL21(DE3). A starter culture (4 mL LB medium supplemented with kanamycin, 50 µg/mL) was grown aerobically overnight at 37˚C with shaking (250 rpm) and used to inoculate 400 mL of LB medium containing kanamycin. Culture was grown at 37˚C with shaking to an OD600 of 0.6 at which point it was induced with IPTG (100 µM final concentration) and incubated for an additional 16 h at 18˚C. Cells were harvested by centrifugation (5,000 × g, 30 min, 4˚C) and resuspended in 25 mL of lysis buffer (20 mM Tris-HCl, 300 mM NaCl, 10 mM imidazole, 5% glycerol, 100 µM phenylmethylsulfonyl fluoride (PMSF)). Cells were lysed by sonication on ice (30 s on, 30 s off cycles, 15 min total), and cell debris was removed by centrifugation (19,000 × g, 30 min, 4˚C). The clarified lysate was applied to a Ni-NTA affinity column containing 0.5 mL of resin pre-equilibrated with equilibration buffer (20 mM Tris-HCl, 300 mM NaCl, and 10 mM imidazole). The resin was washed with 20 column volumes (CV) of wash buffer (20 mM Tris-HCl, 300 mM NaCl, and 25 mM imidazole), followed by sequential washes with 1 CV of 50 mM imidazole and 2 CV each of 100, 150, and 200 mM imidazole. Bound protein was subsequently eluted with 2 CV of 250 mM imidazole. All wash and elution buffers contained 20 mM Tris-HCl and 300 mM NaCl. The eluted protein was concentrated using a 10 kDa molecular weight cutoff centrifugal filter, buffer-exchanged into PBS, and quantified using the Bio-Rad DC Protein Assay.

**In vitro Enzymatic Assays for *N*-butyrylglycine Biosynthesis**

ACOP_v1_740045 activity assays were performed in 20 mM $NH_4HCO_3$ buffer (pH 7.8) with 20 µM purified enzyme, 0.5 mM butyryl-CoA, and 10 mM glycine. Reactions were incubated at room temperature for 1 h and quenched by addition of 50 µL of acetonitrile. Samples were centrifuged (17,000 × g, 5 min) and 20 µL of supernatant was subjected to 3-NPH derivatization. The resulting solution was analyzed by LC-MS as previously described.

**Steady-State Kinetic Analysis of ACOP_v1_740045**

Steady-state kinetic parameters for ACOP_v1_740045 were determined using an Ellman's reagent-based colorimetric assay to quantify the release of free coenzyme A (CoA) during glycine acylation. Enzymatic reactions (15 µL) containing 0.5 µM purified ACOP_v1_740045, 2 mM acetyl-CoA or butyryl-CoA, and varying concentrations of glycine in reaction buffer (50 mM Tris-HCl, pH 8.0) were incubated for 90 sec at 37 °C. Reactions were quenched by adding 15 µL of acetonitrile, followed by centrifugation (16,000 × g, 3 min) to remove precipitated protein. The resulting supernatant was mixed with 90 µL of Ellman's reagent (200 µM 5,5′-dithiobis(2-nitrobenzoic acid), DTNB), and color was allowed to develop for 2 min before measuring absorbance at 412 nm. Apparent Michaelis-Menten parameters were determined by nonlinear regression of reaction velocities against glycine concentration using GraphPad Prism.

**Identification of ACOP_v1_740045 Homologs in Microbial Genomes**

Predicted protein sequences from Human Microbiome Project (HMP) reference genome annotation files (GALES 2018 release) were downloaded from the HMP Data Coordination Center. Protein sequences were combined into a single FASTA database and indexed using NCBI BLAST+. Sequence homology searches were performed using BLASTp with the amino acid sequence of the *N*-acyltransferase ACOP_v1_740045 as the query and an E-value cutoff of 1e-5. For each identified homolog, amino acid identity, alignment coordinates, protein length, E-value, and bit score were extracted. Query coverage was calculated as the number of amino acid positions spanned by the alignment divided by the total query length, expressed as a percentage. Genome accessions were used to retrieve the corresponding bacterial species and strain designations from the NCBI nucleotide database.

Additional homology searches were performed against the Unified Human Gastrointestinal Protein Catalog v2.0 clustered at 90% amino acid sequence identity (UHGP-90). An E-value cutoff of 1e-5 was used. For each identified homolog, amino acid identity, alignment length, query coverage, E-value, and bit score were extracted. Protein identifiers were used to retrieve the corresponding genome accessions for subsequent taxonomic identification.

**Identification of ACOP_v1_740045 Homologs in the Mouse Gut Microbiome**

To investigate the distribution of homologs of the putative biosynthetic enzyme within the mouse gut microbiome, the protein sequence was queried against the MGnify Mouse Gut Protein Catalogue v1.0, containing protein sequences clustered at 95% amino acid identity. A local protein BLAST database was constructed from the catalogue FASTA file and sequence similarity searches were performed using BLASTP with an E-value threshold 1e-5. Protein homologs were evaluated based on amino acid sequence identity, query coverage, and E-value. Taxonomic assignments were obtained by cross-referencing the genome accession associated with each BLAST hit against the MGnify Mouse Gut Genome Catalogue metadata.

**Cell-Cycle Analysis by Propidium Iodide Staining and Flow Cytometry**

Caco-2 cells were seeded at a density of $1 \times 10^5$ cells/well in 24-well tissue culture plates and allowed to adhere for 24 h at 37 °C and 5% $CO_2$. Cells were subsequently treated with 0.5 mM *N*-butyrylglycine or vehicle control for 6 h. Following treatment, culture medium was removed, and cells were washed once with PBS before detachment with 0.05% trypsin (3 min, 37 °C). Trypsinization was quenched with fresh culture medium, and cells were collected by centrifugation (350 × g, 3 min). Cell pellets were washed twice with PBS and resuspended in 100 μL of PBS. Ice-cold 70% ethanol (1 mL) was added with gentle mixing, and cells were fixed at −20 °C for 24 h.

Following fixation, cells were collected by centrifugation (350 × g, 3 min), washed twice with cold PBS, and pelleted. Cell pellets were resuspended in 50 μL of RNase A (100 μg/mL) and 400 μL of propidium iodide (PI; 50 μg/mL). Samples were gently mixed and incubated for 30 min at room temperature protected from light. Cellular DNA content was analyzed using a BD Accuri C6 Plus flow cytometer (BD Biosciences), and data were processed using BD Accuri software. Cell aggregates and doublets were excluded by gating on PI fluorescence area versus width (PI-A vs. PI-W). DNA-content distributions were evaluated using PI fluorescence histograms.

#### Primers and Constructs

>ACOP_v1_740045 Construct (codon optimized for *E. coli*)

ATGATTACCCAAGTACGTACACCCCGTCAGCGTCAACGATTTCGTTCCGCCTGTCGTGGTAAACT
CTGTTTGGGTGTGACTATGCCCCAAGCTTTGGCTTTGTTCGGCAAGAGCCAACCGGGTCGGTTCTT
CGCTGGTCCTACACTTGCGCTGGACGTGGGCGGGTCAACCGCCTGGTTGGCTGGTCACGCAAACC
CCGATGAGTTAGCGAGTTTTCTGCATTTCTGCGGCTGCAAGGCTGTAATTCTTGACGAGGCAGAG
TGCCCGCCCCCAACTGGGTGGGCTCGGGCCAAGAGCCACTCAGTATTTGGTCTTGCACCTGGCAA
GCAACTTCCAGAGCCGGCTGTCGAGGAAACACTTTGGGCATCTCTGCACTTTGACCCCGAGCCAC
CCGCGGGCCCTGTTGCACAGATGTTATTCCCAGATCGCCCGACAAGACGAGACGACTTCTATTCT
GAACTCTGCAGTAAGCGCGCGCGTGGCCGAGCAGTCGTCTGGGCGTTAGAGCAAGGTGGGGAGT
TAGCCTGCACGGTAGGTGCGTACGCGATTGGTACCGAGCAAGCTTACATGGCGTGCGGGGAAAC
GGCAGAGCCGTTAAGAGGTAGAGGCATAGGTGGGCGGCTGATTGTCCGCTTGGCTAATACCTTG
AGTGCCCAGGGCTTACAACCCCTGTTCCTTTGTTCACCGGAGCGCGTGCATTTCTACACCAGATTG
GGTTTTGAAAAGCTCGGGGAGTACGCCCGTTACGAGAATAAGGATCAGGGA

| **Primers** | Sequence 5’ -> 3’ |
|---|---|
| RPLP0 Forward qRT-PCR Primer | TGGTCATCCAGCAGGTGTTCGA |
| RPLP0 Reverse qRT-PCR Primer | ACAGACACTGGCAACATTGCGG |
| MKI67 Forward qRT-PCR Primer | CTTTGGGTGCGACTTGACGA |
| MKI67 Reverse qRT-PCR Primer | ACAACTCTTCCACTGGGACG |
| AOC1 Forward qRT-PCR Primer | AGCAGAGCACACAGAGCC |
| AOC1 Reverse qRT-PCR Primer | GCCTTCCTGGGCAGAGTC |

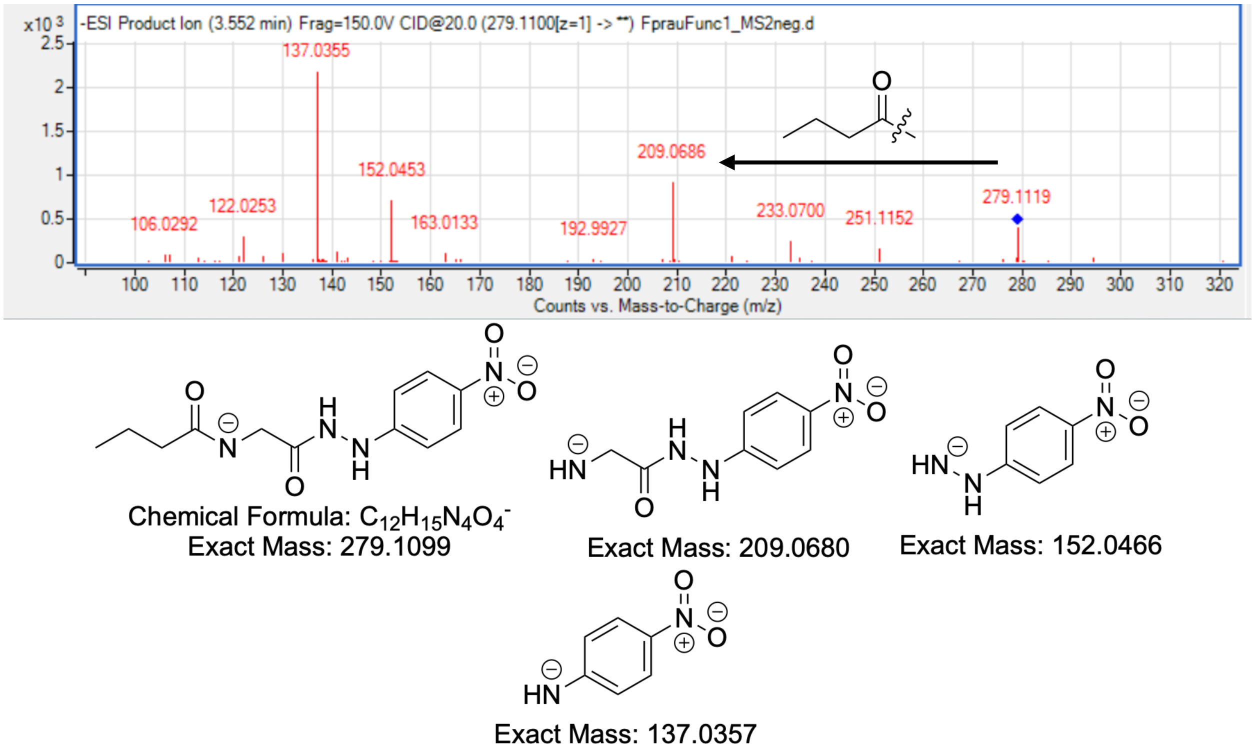


**Figure S1**. ESI$^-$-HRMS/MS fragmentation spectrum of natural 3-NPH-derivatized *N*-butyrylglycine (20 eV). Structures consistent with fragmentation pattern are shown.

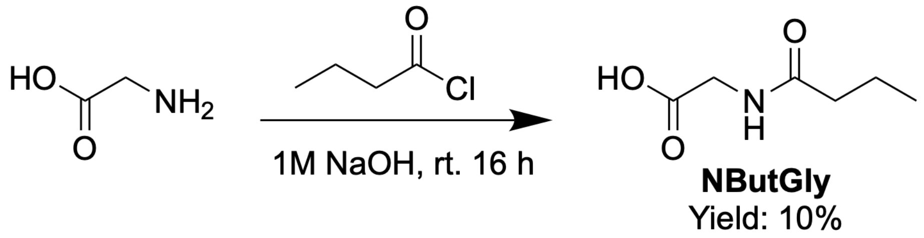


**Figure S2**. Synthesis of *N*-butyrylglycine.

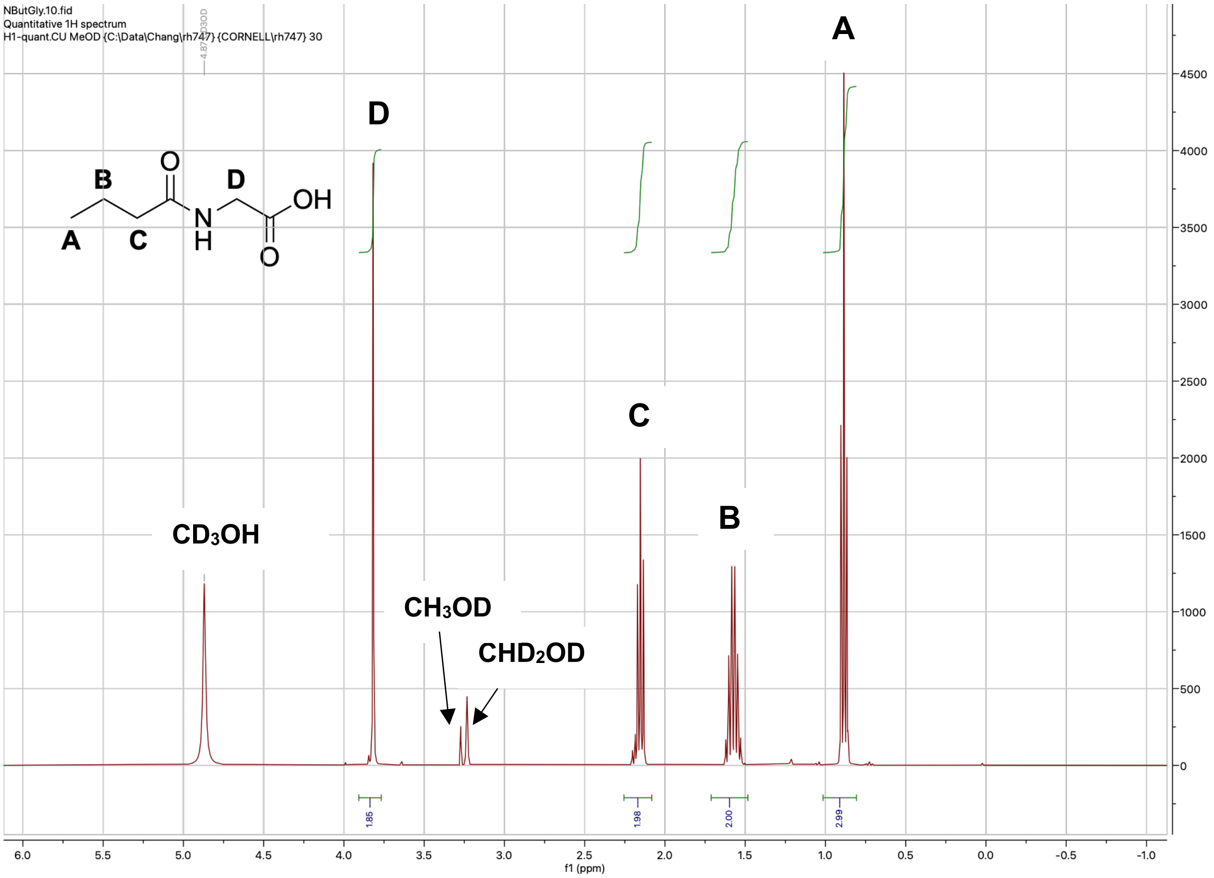


**Figure S3**. [1]H NMR spectrum of *N*-butyrylglycine (400 MHz, methanol-$d_4$).

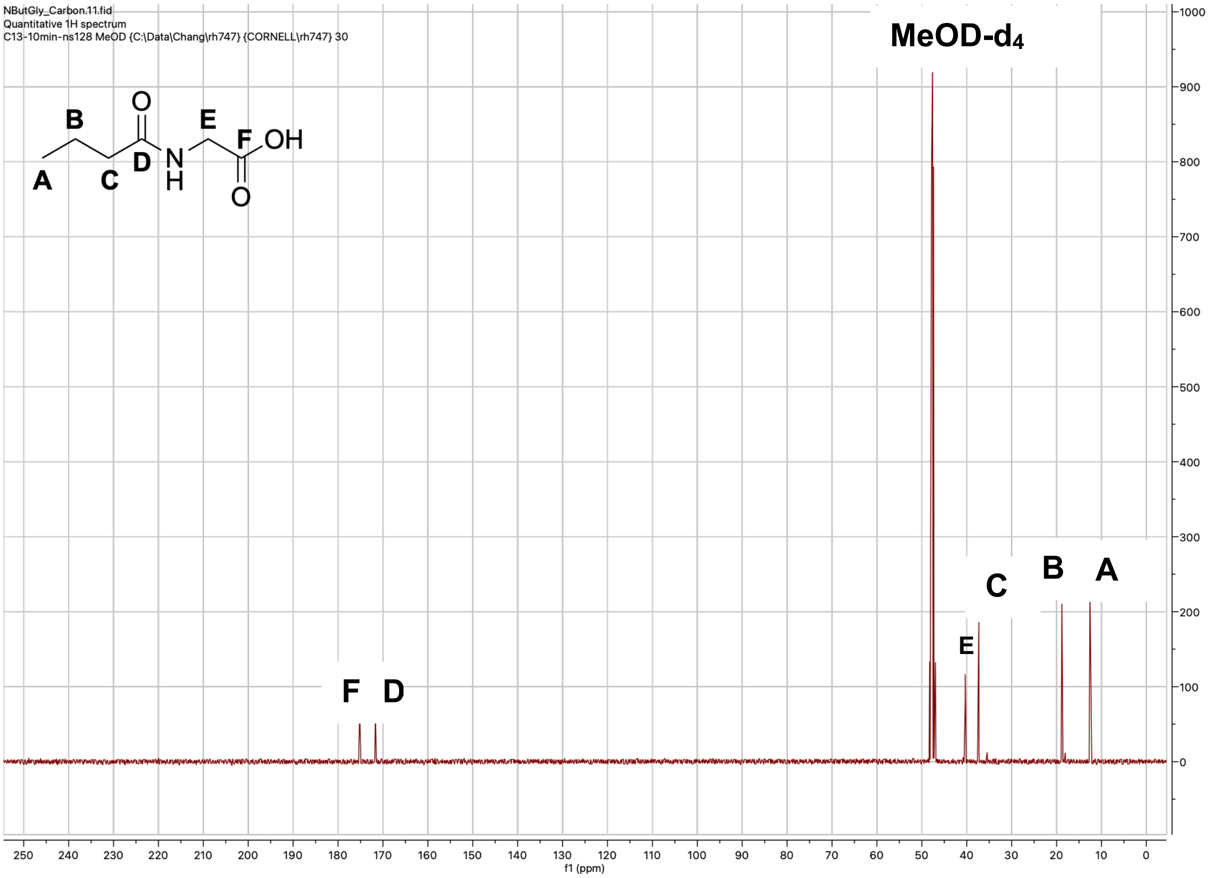


**Figure S4**. $^{13}$C NMR spectrum of *N*-butyrylglycine (400 MHz, methanol-$d_4$).

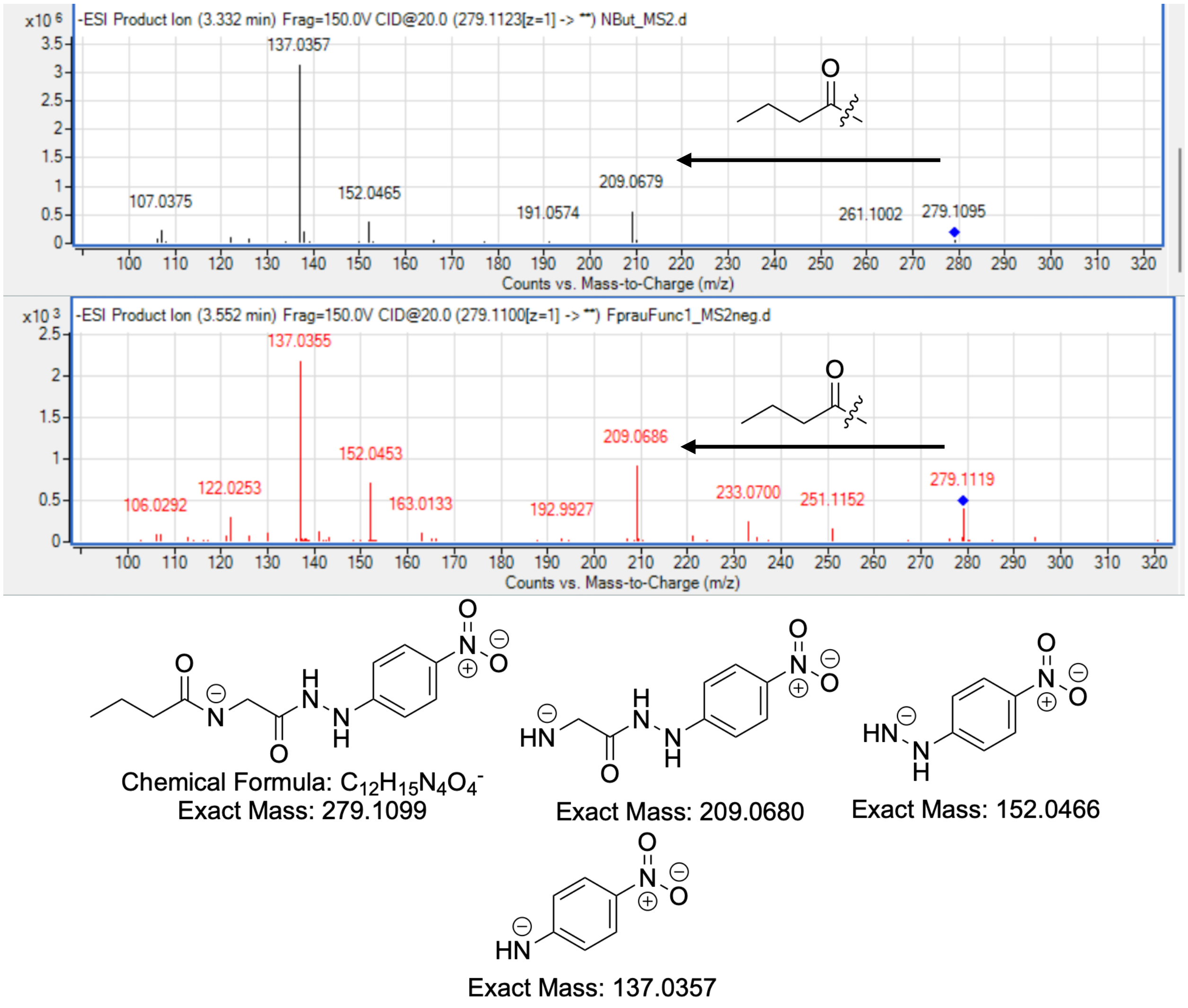


**Figure S5**. ESI$^-$-HRMS/MS fragmentation spectrum of synthetic (top) and natural (bottom) 3-NPH-derivatized *N*-butyrylglycine (20 eV). Structures consistent with fragmentation pattern are shown.

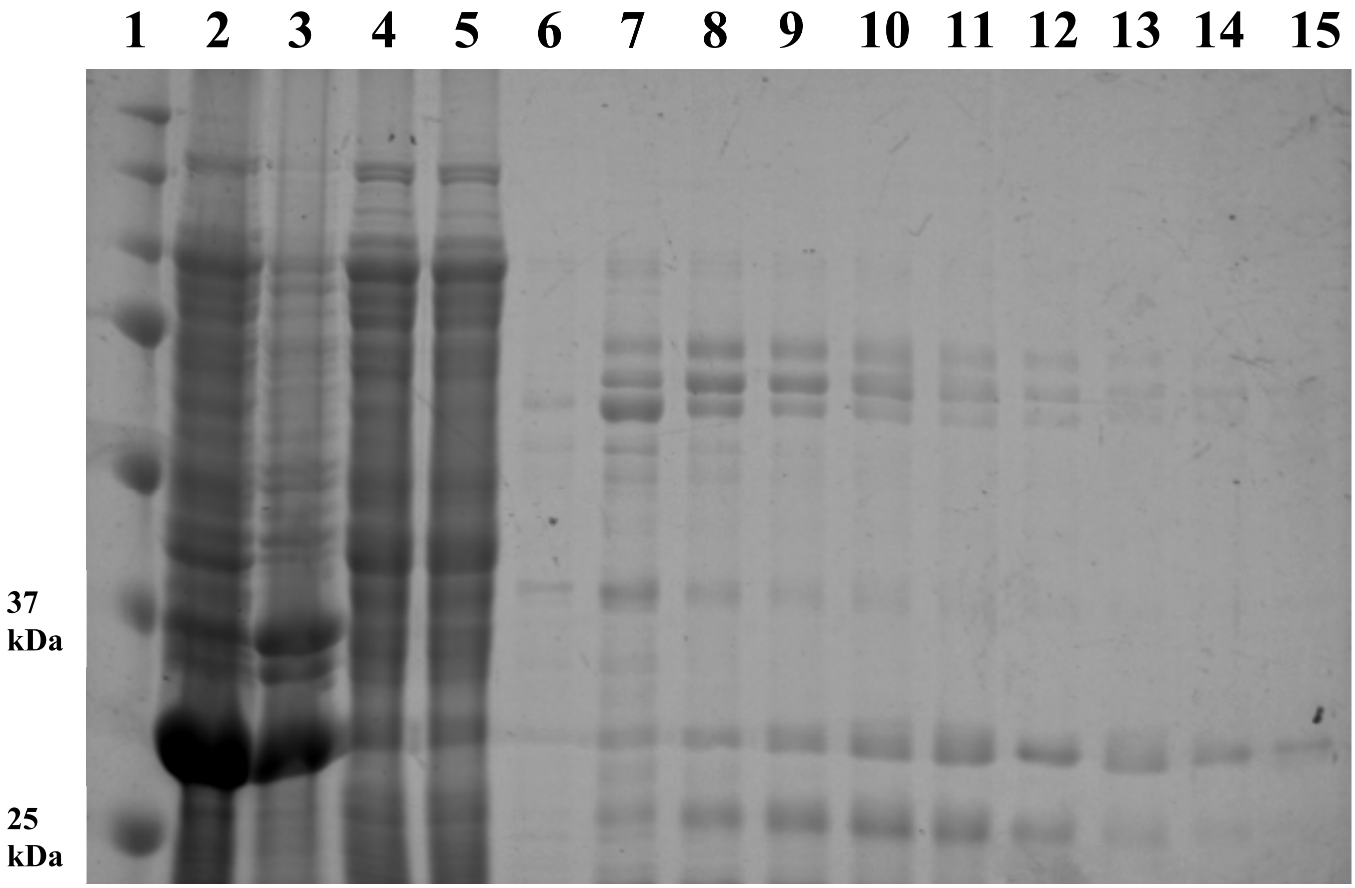


**Figure S6**. **Purification of ACOP_v1_740045**. Recombinant ACOP_v1_740045 (31 kDa) was analyzed by 12% SDS-PAGE. Electrophoresis was performed at 100 V for 10 min, followed by 180 V for 1 h. Fractions corresponding to lanes 14 and 15 were pooled and concentrated using a 10 kDa molecular weight cutoff (MWCO) Amicon Ultra centrifugal filter. Lane identities are listed below.

**1: Ladder**
**2: Lysate**
**3: Pellet**
**4: Flowthrough**
**5: 25 mM Imidazole Wash 1**
**6: 25 mM Imidazole Wash 20**
**7: 50 mM Imidazole Wash 1**
**8: 100 mM Imidazole Wash 1**
**9: 100 mM Imidazole Wash 2**
**10: 150 mM Imidazole Wash 1**
**11: 150 mM Imidazole Wash 2**
**12: 200 mM Imidazole Wash 1**
**13: 200 mM Imidazole Wash 2**
**14: 250 mM Imidazole Elute 1**
**15: 250 mM Imidazole Elute 2**

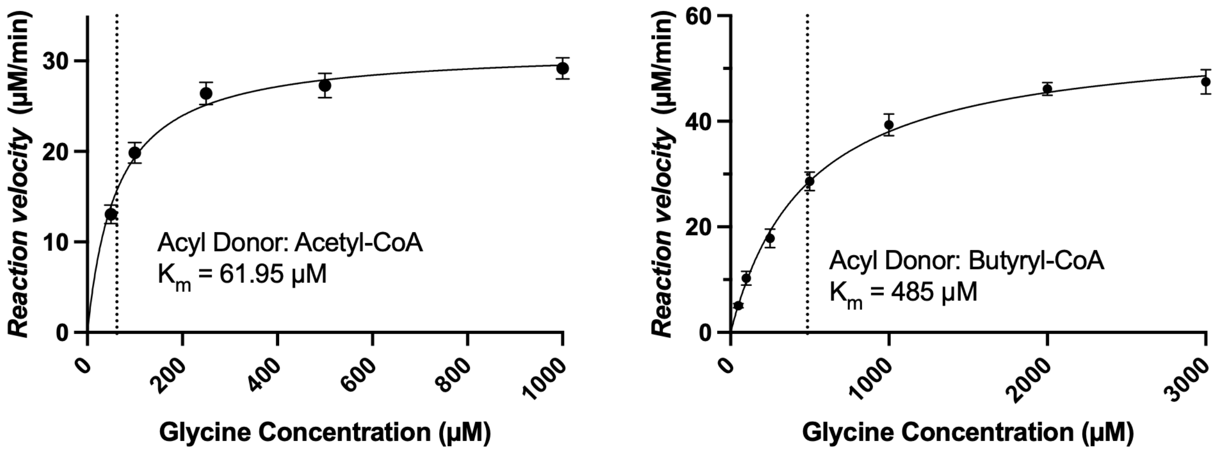


**Figure S7**. **Steady-state kinetic characterization of ACOP_v1_740045-mediated glycine acylation**. Michaelis-Menten plots showing reaction velocity as a function of glycine concentration using acetyl-CoA (left) or butyryl-CoA (right) as the acyl donor. Reaction velocities were determined using an Ellman's reagent-based colorimetric assay to monitor CoA release. Apparent $K_m$ values for glycine were 61.95 µM with acetyl-CoA and 485 µM with butyryl-CoA. Dotted vertical lines indicate the corresponding apparent $K_m$ values. Apparent $k_{cat}$ values for glycine were 1.05 $s^{-1}$ with acetyl-CoA and 1.88 $s^{-1}$ with butyryl-CoA. Acyl-CoA concentrations were held constant at 2 mM. Data are presented as mean ± SD (n = 3).

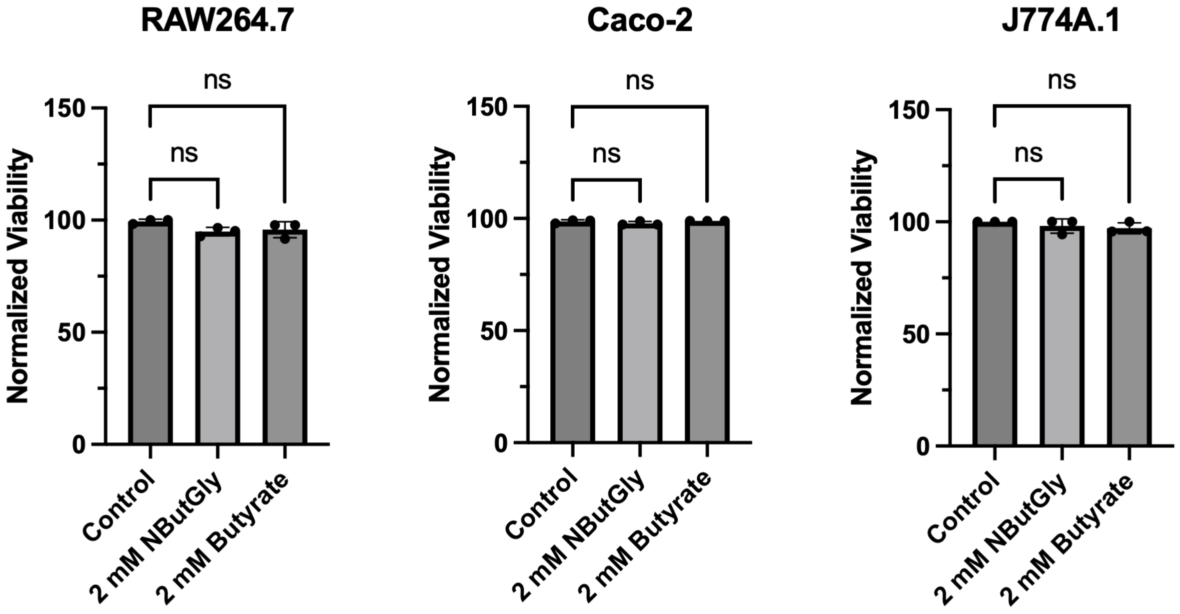


**Figure S8**. **Effects of NButGly and butyrate treatment on mammalian cell viability**. Cell viability was assessed by trypan blue exclusion in RAW 264.7, Caco-2, and J774A.1 cells following treatment with 2 mM NButGly or butyrate. Viability was normalized to untreated controls. Neither treatment significantly affected cell viability relative to control in any of the cell lines examined. Data are presented as mean ± SD. Statistical significance was determined using Welch's t-test; not significant (ns). All data represent at least three biological replicates.

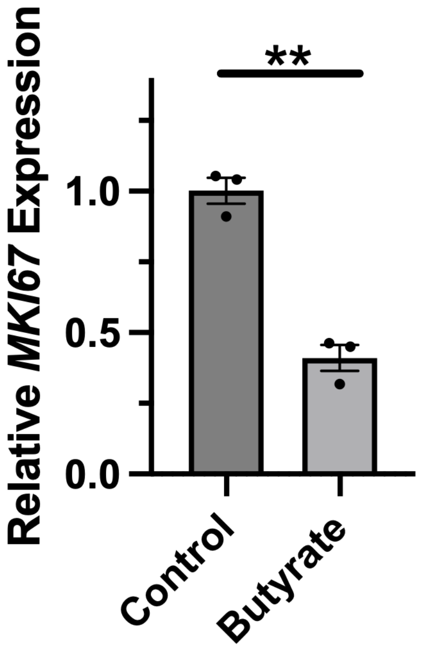


**Figure S9**. **Effect of butyrate treatment on *MKI67* expression in Caco-2 cells**. Relative *MKI67* mRNA expression was measured by qRT-PCR following treatment (6 h) with butyrate (0.5 mM) and normalized to untreated controls. Data are presented as mean ± SEM. Statistical significance was determined using Welch's t-test; $P \leq 0.01$ (**). All data represent at least three biological replicates.

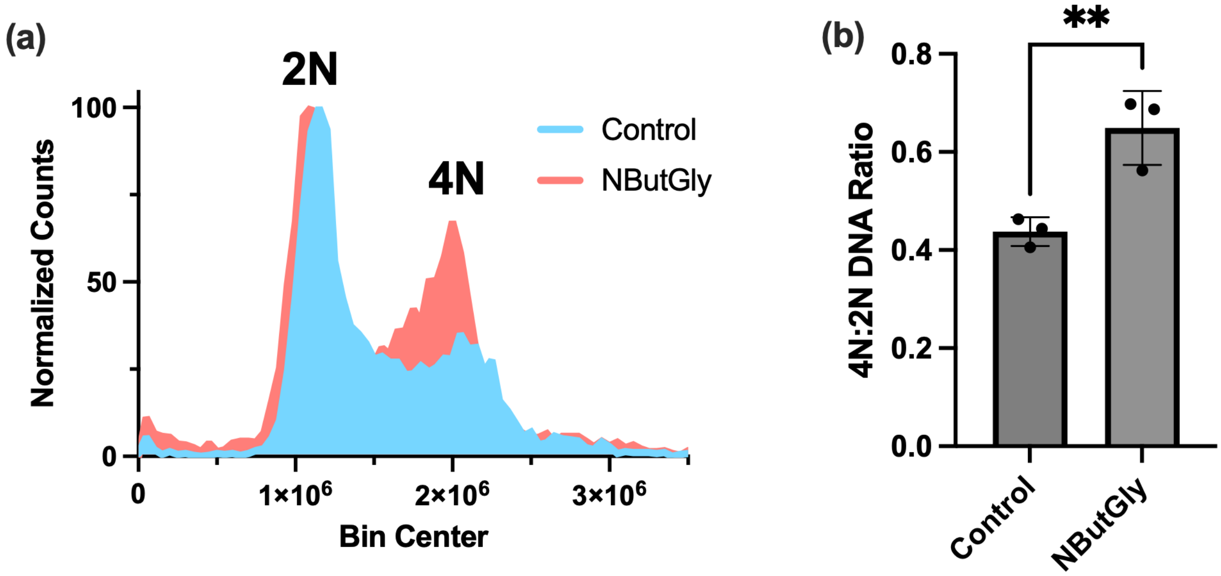


**Figure S10**. **Effect of NButGly treatment on cellular DNA-content distribution in Caco-2 cells**. (a) Representative propidium iodide (PI) staining profiles obtained by flow cytometry showing the distribution of cellular DNA content in control and NButGly-treated Caco-2 cells. Peaks corresponding to 2N and 4N DNA content are indicated. (b) Quantification of the 4N-to-2N ratio, showing an increased relative 4N DNA-content signal following NButGly treatment. Data are presented as mean ± SD. Statistical significance was determined using Welch's t-test; $P \leq 0.01$ (**). All data represent at least three biological replicates.

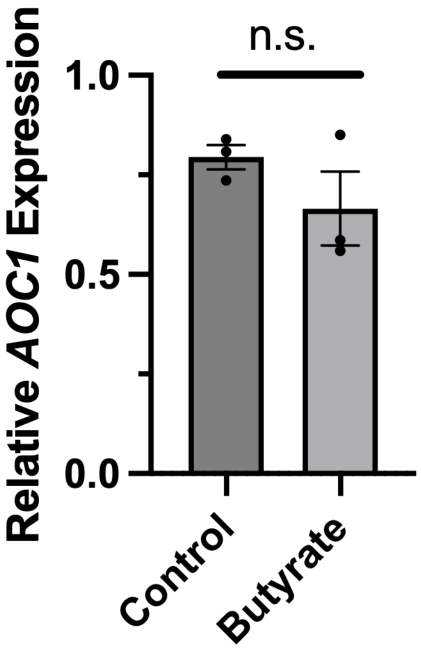


**Figure S11**. **Effect of butyrate treatment on *AOC1* expression in Caco-2 cells**. Relative *AOC1* mRNA expression was measured by qRT-PCR following treatment (6 h) with butyrate (0.5 mM) and normalized to untreated controls. Data are presented as mean ± SEM. Statistical significance was determined using Welch's t-test; not significant (n.s.). All data represent at least three biological replicates.
